# Human newborns form musical predictions based on rhythmic but not melodic structure

**DOI:** 10.1101/2025.02.19.639016

**Authors:** R. Bianco, B. Tóth, F. Bigand, T. Nguyen, I. Sziller, G.P. Háden, I. Winkler, G. Novembre

**Author notes:** **Correspondence:** Roberta Bianco, Neuroscience of Perception & Action Lab, Italian Institute of Technology, Viale Regina Elena 291, 00161, Rome, Italy.

## Abstract

The ability to anticipate rhythmic and melodic structures in music is considered a fundamental human trait, present across all cultures and predating linguistic comprehension in human development. Yet, it remains unclear the extent to which this ability is already developed at birth.

Here, we used temporal response functions to assess rhythmic and melodic neural encoding in newborns (N = 49) exposed to classical monophonic musical pieces (real condition) and control stimuli with shuffled tones and inter-onset intervals (shuffled condition). We computationally quantified context-based rhythmic and melodic expectations and dissociated these high-level processes from low-level acoustic tracking, such as local changes in timing and pitch.

We observed encoding of probabilistic *rhythmic* expectations only in response to real but not shuffled music. This proves newborns’ ability to rely on rhythmic statistical regularities to generate musical expectations. We found no evidence for the tracking of *melodic* information demonstrating a downweighing of this dimension compared to the rhythmic one.

This study provides neurophysiological evidence that the capacity to track statistical regularities in music is present at birth and driven by rhythm. Melodic tracking, in contrast, may receive more weight through development with exposure to signals relevant to communication, such as speech and music.

**Significance:** Perception and appreciation of music stem from humans’ universal ability to track sequential pitch and timing variations, forming musical melodies and rhythms. To investigate the neurobiological origins of this capacity, we measured neural encoding of rhythm and melody in newborns listening to naturalistic music. We demonstrate a precocious ability to track high-level rhythmic statistical regularities beyond low-level acoustic features. Conversely, melodic tracking is virtually absent at birth, likely emerging with later exposure to pitch-varying signals such as speech and music. This reveals that newborns gradually rely on different sound dimensions to start making sense of the auditory environment. While rhythm and melody are both universal components of music, they do not emerge in parallel but instead develop along distinct timelines.

## Introduction

Music is an increasingly compelling means for understanding the development of a wealth of neuro-cognitive processes, including those that support communication through sound patterns (1). Already at the earliest stages of development, the human brain draws on multiple auditory cues to extract meaningful patterns, such as words or melodies, from the acoustic environment (2, 3). This process is facilitated by the integration of sequential information and, thereby, by the extraction of statistical patterns along temporal and spectral dimensions, such as timing and pitch (4, 5). In music, tracking of statistical patterns is largely implicit (6), and it allows the brain to link each note with past events, namely its rhythmic (temporal) and melodic (spectral) structural context. By drawing on past contexts, listeners can recognize patterns (7) and leverage this information to anticipate *when* an event will occur and *what* it will be (8–10). Such rhythmic and melodic expectations are the backbone of music perception and appreciation (11) and are assumed to have contributed to the evolution and development of human musicality (12–17).

Based on cross-species studies, rhythmic and melodic expectations in primate species seem to have evolved along different phylogenetic pathways. Sensitivity to rhythmic patterns was observed in non-human primates, suggesting deep phylogenetic roots (18–23). In contrast, the sensitivity to melodic patterns based on pitch relations appears more variable, if not absent, in non-human primates and may be unique to humans within the primate lineage (18, 24–26). This observation raises an important question: are humans naturally predisposed to melodic tracking? Answering this question is challenging yet important for understanding how innate traits and cultural influences shape the complex spectrum of human musical abilities observed worldwide (27, 28).

Here, we take human newborns as a testbed for studying the human brain’s predisposition to process music, specifically its rhythmic and melodic aspects. Newborns’ auditory responses can be reliably recorded using electroencephalography (EEG) (29, 30), and these responses are marginally influenced by prior exposure compared with those measured at any later developmental stage (but see Giordano et al., 2021; Keller, 2012; Lang et al., 2020; Moon et al., 1993; for a review James, 2010). Compelling evidence suggests that the human brain engages with sounds already in utero, as fetuses discriminate, habituate to, and memorize sounds (36). By 28-35 weeks of gestation, fetuses can differentiate between music and speech, showing increased heart rates and body movements in response to music (37). In terms of rhythm perception, EEG studies demonstrated that newborns already possess some rhythmic capacities, such as adaptation to the presentation rate of temporal patterns (38), tracking of meter-related frequencies (39), and perception of the beat (40). What remains unclear is whether newborns can use rhythmic statistical regularities beyond sound periodicities, such as transition probabilities to form temporal expectations (41). In terms of melodic capacities, EEG studies showed that newborns exhibit discrimination of pitch independent of timbre and detection of highly surprising events, such as deviants from deterministic patterns of tones or regularities in sequences of tone intervals (44, 45). These studies provide preliminary evidence for expectations based on probabilistic distributions of melodic information. Yet they tested only the two tail-ends of such presumed probabilistic distribution: very frequent vs. very infrequent events, ignoring the wide range of note-by-note surprises of real music. This leaves it unclear whether newborns can form melodic expectations whilst listening to continuous natural music, as observed in adults (18, 46, 47). Finally, because melodic and rhythmic abilities have often been studied separately, the weights of rhythmic and melodic expectations during music processing at birth are unknown. Here, we investigate neural tracking of probabilistic expectations based on both timing and pitch structures to understand how the newborn brain weights these musical features while listening to ecologically valid musical stimuli.

Rhythmic and melodic expectations can be generated through different anticipatory mechanisms sensitive to different features of the stimulus – from surface acoustical attributes to local and global event-based statistics. Thus, using the multivariate Temporal Response Function analysis (mTRF) (48, 49), we measured how multiple features of the continuous musical stimuli – ‘low-level’ acoustic features and ‘high-level’ probabilistic rhythmic/melodic information – predict neural EEG responses of human newborns. As in previous human and non-human primate work (18, 46, 47), we assessed neural encoding of J. S. Bach’s piano monophonic pieces – rich musical stimuli combining both melodic and rhythmic probabilistic structures. Based on previous findings of rhythmic but not melodic tracking in non-human primates (18), we hypothesized that human newborns would show a similar pattern if these abilities were inherited phylogenetically. This would imply that whilst rhythm encoding is embedded in the human brain from the outset, melodic encoding might develop more slowly with experience and behavioral relevance. Conversely, if, unlike other non-human primates, rhythmic and melodic sensitivity emerges in parallel in humans, then human newborns might already exhibit some capacity for melodic encoding, potentially comparable to rhythmic encoding, as observed in adults (18, 46).

## Results

A mTRF analysis was carried out to assess the neural encoding of musical expectations at birth (Fig. 1A). Newborns were exposed to musical melodies (real condition) and control stimuli (shuffled condition, where pitch and note timings were shuffled over time to create sequences with disrupted musical regularities). Probabilistic expectations were estimated based on the information-theoretic properties of the stimuli using a variable-order Markov model of statistical learning (i.e., information dynamic of music [IDyOM]; (50)). The model leverages observations from the past (long- and short-term) musical context to compute Shannon’s surprise (S) and entropy (E) of each note of each melody, specifically concerning pitch (Sp and Ep, respectively) and onset timing (St and Et). According to these estimates, shuffled melodies were overall more unexpected than real melodies, both for pitch (Sp: W = 40, p = .002; Ep: W = 40, p = .002; see Methods ‘Statistical analysis’ for details) and timing (St: W = 35, p = .036; Ep: W = 33, p = .075) (Fig. 1B).

**Figure 1.**
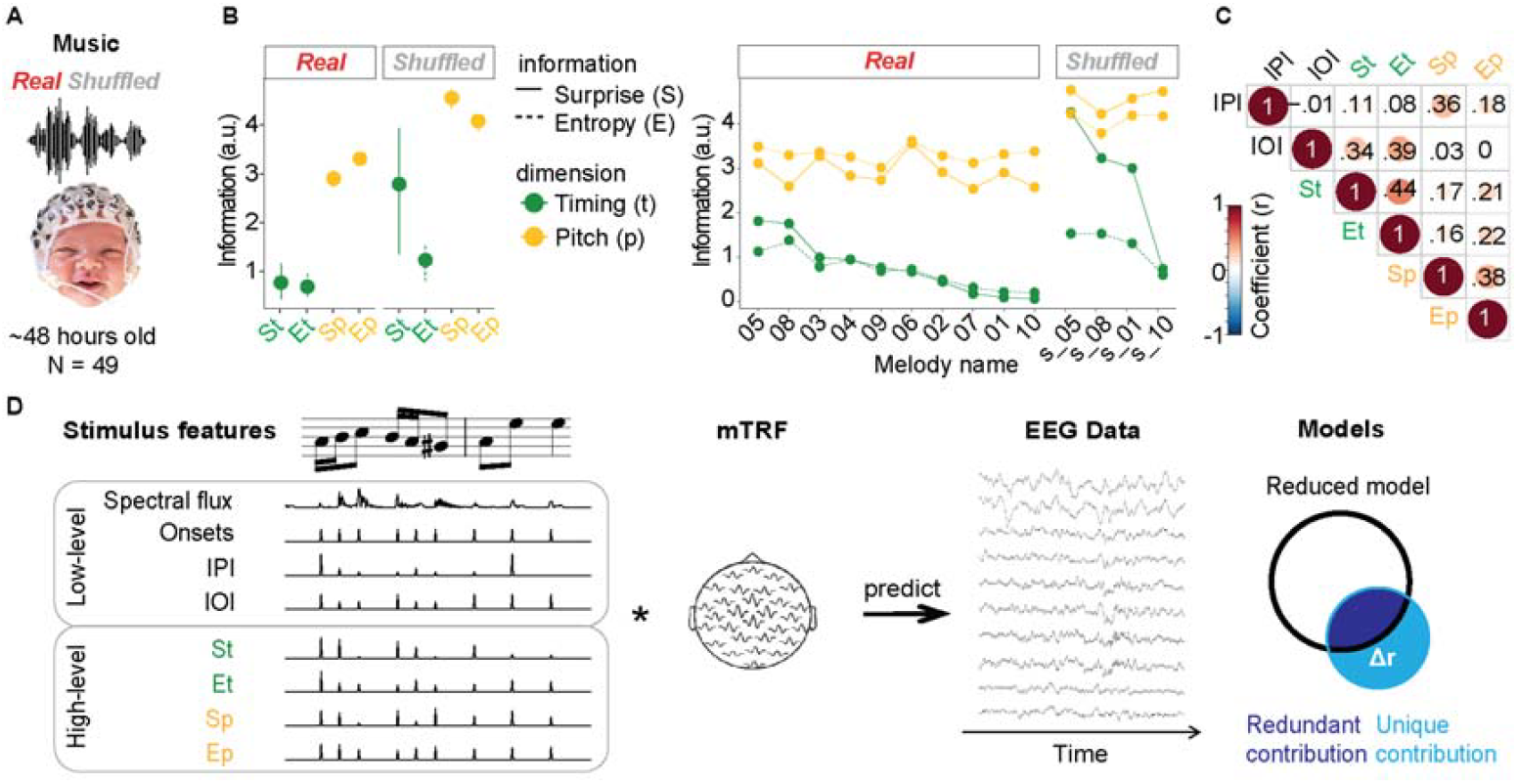
Materials and Methods. A) Experimental paradigm. We analyzed EEG data recorded from 49 sleeping human newborns while being exposed to monophonic piano melodies composed by J. S. Bach (Real condition) and control stimuli (Shuffled condition). **B) Surprise and entropy**. Surprise and entropy associated with each note’s timing (green, St and Et, respectively) and pitch (yellow, Sp and Ep) were estimated using an unsupervised statistical learning model trained on all stimuli. Dot plots display mean surprise and entropy associated with real and shuffled music, averaged across melodies (left panel), and separately for each melody (right panel). Error bars represent bootstrapped 95% confidence intervals (CI). **C) Correlations between stimulus features**. Pearson’s correlations (r values) between the stimulus features: inter-pitch-interval (IPI), inter-onset-interval (IOI), and Surprise and Entropy associated with timing (St and Et) and pitch (Sp and Ep). **D) Analytical approach**. Multivariate Temporal Response Function (mTRF) models were fit to describe the forward relationship between multiple stimulus features and the EEG signal. The full TRF model (leftmost panel) included acoustic low-level features (Spectral flux, Acoustic Onset, IOI, IPI) and high-level features (Surprise and Entropy of pitch and timing). To assess the unique contribution of each feature (or set of features) to the EEG data, we run reduced models encompassing all variables but with the specified one being randomized in time (yet preserving the note onset times). We then calculated the difference in EEG prediction accuracy (Pearson’s correlations, r) between the reduced models and the full model (Δr). On the rightmost panel, the black circle denotes information of a reduced model, with the variable of interest being randomized. The single variable (blue circle) explains redundant information (dark blue) with the reduced model and unique information (light blue) that increases the explanatory power of the full model (that includes the area of both the black and the light blue circles).

Each note’s surprise and entropy estimate positively correlated with low-level acoustic features values, such as the magnitude of the latest pitch or timing interval (i.e., inter-pitch-interval, IPI, or inter-onset-interval, IOI) (Fig. 1C). Hence, we assessed the unique contribution of probabilistic pitch and timing expectations to the EEG data, in addition to the contribution from low-level acoustic processing (including IPI, IOI, acoustic onsets and spectral flux) (Fig. 1D). To do so, we derived single-participant TRFs by fitting multivariate lagged regression models (Fig. 2A). We then estimated prediction accuracy (Pearson’s correlations, *r*) between the EEG signals predicted by the TRF models and the actual EEG data (averaged across all participants, ‘ground-truth EEG data’ see Methods ‘TRF analysis’), separately for each melody and EEG channel, using leave-one-melody-out cross-validation over a lag window ranging from -50 to 400 ms. To assess the unique contributions of the variables of interest to the EEG data, we trained reduced models that included a veridical representation of all variables except for the variable of interest, which was randomized (see methods TRF Analysis). Finally, we calculated the difference in prediction accuracy (Δr) between the reduced and full models, selecting the top 25% of channels with the highest prediction accuracy in the full model across conditions.

**Figure 2.**
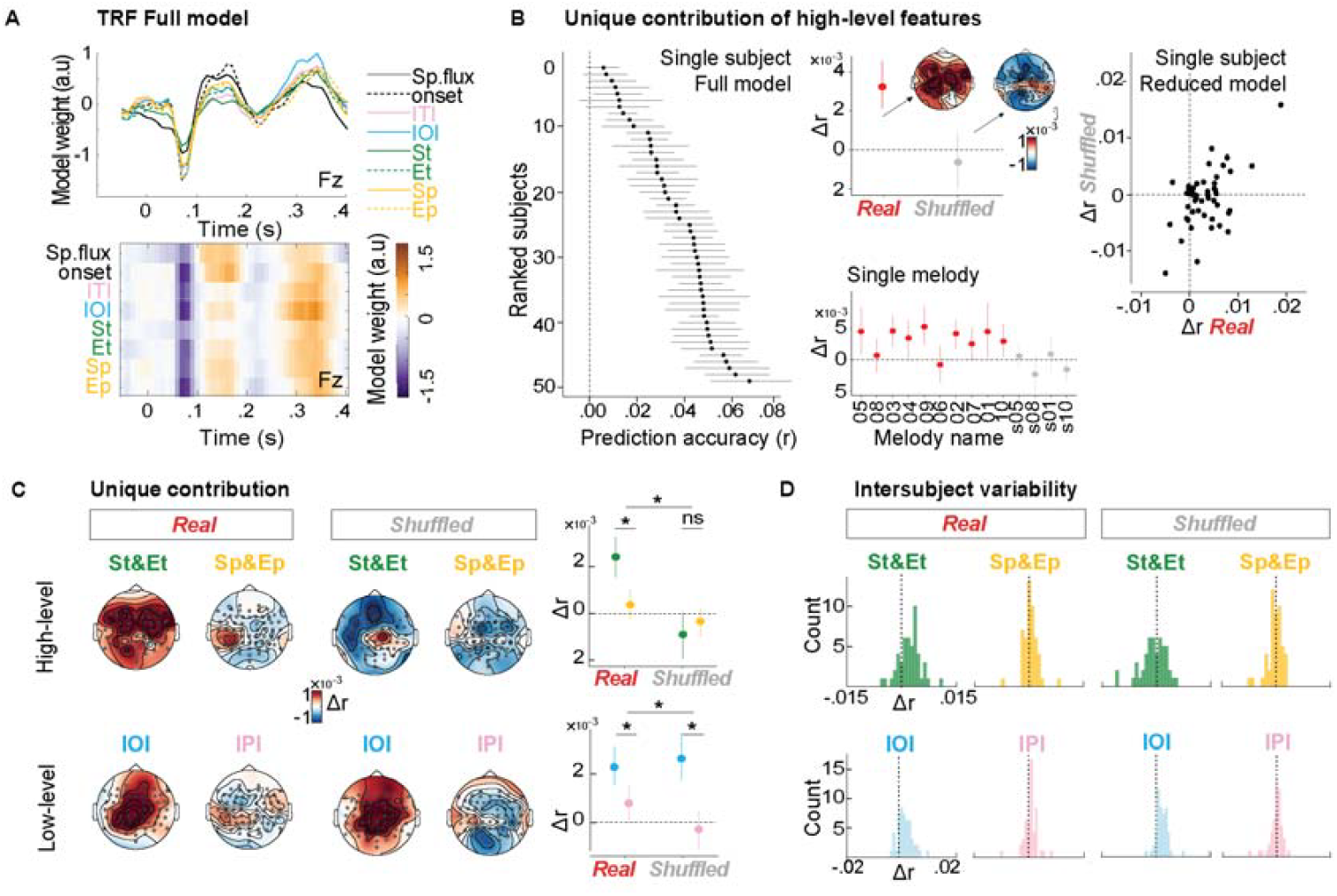
Neural encoding of timing but not pitch expectations at birth. A) TRF full model. Ridge regression weights (see Methods ‘TRF analysis’) in time yielded by TRF for all predictors of the full model at the electrode Fz. Orange and purple colors indicate positive and negative weights, respectively. Zero is at the note onset. **B) Unique contribution of high-level musical features**. Left panel: black dots indicate EEG prediction accuracy of the full model of each infant, computed across 25% of channels with the highest prediction accuracy across conditions (best channels per infant, used in the following plots). Infants are ranked according to the prediction accuracy (r) of the full model. Central panels: group-average (n=49) Δr resulting from the difference between full and reduced models assessing the unique contribution of high-level musical features (St and Et + Sp and Ep) computed across each infant’s best channels and plotted separately for real (red) and shuffled (grey) melodies (lower panel; the associated topographical maps are shown on the upper panel. Error bars represent bootstrapped 95% CI. Right panel: scatter plot representing the relationship between Δr yielded by high-level musical features associated with real (x-axis) and shuffled (y-axis) melodies (each participant is represented by a dot). **C) Unique contribution of timing and pitch-related features**. Left: Topographical maps representing group-average Δr resulting from the difference between full and reduced models separately assessing the unique contribution of St and Et, Sp and Ep, IOI, and IPI to the models of EEG data across conditions. Right: dot plot representing the group-average mean Δr’s for the four reduced models, separately for the two conditions (dot colors match those on the left panel), computed across each infant’s best channels. Error bars represent bootstrapped 95% CI; asterisks indicate the presence of significant main effects and interactions; ‘ns’ indicates non-significant effects. **D) Intersubject variability**. Histograms of individual Δr’s for models reduced by St and Et, Sp and Ep, IOI, and IPI, separately for the two conditions.

### Encoding of probabilistic expectations in real but not shuffled music

Fig 2B (left panel) shows that, despite substantial intersubject variability, the full model, including all features, could predict EEG data with reasonable accuracy across virtually all participants. To what extent do probabilistic (high-level) expectations contribute to the neural signal? We tested the unique contribution of probabilistic (high-level) features derived from the IDyOM model (St and Et + Sp and Ep) beyond low-level stimulus features (Onset, Spectral flux, IOI, IPI). We thus compared the Δ*r* (Full - minus – a reduced model with randomized St and Et + Sp and Ep) across real and shuffled music (Fig. 2B top central panel). A linear mixed effect model (see Methods ‘Statistical analysis’ for details) with the fixed factor Condition (real vs shuffled) yielded a main effect of Condition (χ^2^(1) = 12.065, p < .001) indicating encoding of probabilistic expectations in real but not shuffled music (real > shuffled: b = .004, SE = .001, p = .005; real > 0: b = 0.003, SE = .0007, p < .001; shuffled > 0: p = .594). These effects were not driven by any specific melody (Fig. 2B, bottom central panel) and exhibited high variability across subjects (Fig. 2B, left panel). This analysis demonstrates that the predictable structure of real (but not shuffled) melodies allows babies to generate musical expectations over and above mere acoustic tracking.

### Timing-but not pitch-related expectations

We tested whether the encoding of probabilistic expectations was specifically driven by pitch or timing structures (Fig. 2C). We thus examined the difference between the full model and a reduced model, in which either St and Et (timing probabilistic TRF model) or Sp and Ep (pitch probabilistic TRF model) were randomized. A linear mixed effect model with fixed factor Condition (real vs shuffled) and TRF model (St and Et vs Sp and Ep) yielded a main effect of condition χ^2^(1) = 12.353, p < .001) and an interaction between Condition and TRF model (χ^2^(1) = 9.897, p = .002). For real music, paired contrasts indicated encoding of probabilistic expectations based on timing but not pitch structure (St and Et > Sp and Ep: b = .002, SE = 0.0004, p < .001; St and Et > 0: b = 0.0024, SE = .0005, p < .001; Sp and Ep > 0: p = .384), whereas for shuffled music, neither of the two dimensions yielded significant effects (St and Et > Sp and Ep: p = .856; St and Et > 0: p = .166; Sp and Ep > 0: p = .601). This analysis demonstrates that newborns track the predictable rhythmic structure of the real melodies to generate expectations. In contrast, pitch-based probabilistic expectations do not appear to emerge with statistical significance.

As a control, we ran similar analyses to test the unique contribution of expectations driven by just immediate local changes in timing and pitch, as estimated by IOI and IPI. We thus examined the difference between the full model and a reduced model, in which either IOI or IPI were randomized. A linear mixed effect model with fixed factor Condition (real vs shuffled) and TRF model (IOI vs IPI) yielded a main effect of TRF model χ^2^(1) = 48.225, p < .001) and an interaction between Condition and TRF model (χ^2^(1) = 5.032, p = .025). Paired contrasts indicated encoding of IOI but not IPI for both real (IOI > IPI: b = .001, SE = 0.0003, p = .001; IOI > 0: b = 0.0022, SE = .0005, p < .001; IPI > 0: p = .12) and shuffled music (IOI > IPI: b = .003, SE = 0.0005, p < .001; IOI> 0: b = .00026, SE = .0007, p = .002; IPI > 0: p = .705). This analysis demonstrates that expectations based on local temporal intervals are not altered by the rhythmic structure of the music, as IOIs were similarly tracked in real and shuffled melodies. It also shows that encoding of the pitch information did not reach significance in either condition (although IPI tracking approached significance when compared to zero in the real condition). Hence, the current results do not support the tracking of either probabilistic expectations or local pitch change.

### Event-related potentials (ERPs)

To ground the TRF results in more widely used neurophysiological responses, we examined ERP responses to a subsection of musical notes, i.e. those carrying the highest and lowest 20% quantiles of surprise values (High S and Low S, respectively), separately for pitch and timing (Fig. 3A). The ERPs consisted of a first negative peak (termed N1) followed by two broad positive-going deflections (P1 and P2) separated by a small (second) negative-going deflection (N2). The ERP waveforms resemble previously observed ones evoked by auditory stimuli in newborns (51). Further, the waveform is reminiscent of TRF’s regression weights (Fig. 2A), suggesting that the TRF analysis primarily captured phase-locked auditory responses, as observed in previous studies (18, 46, 52).

**Figure 3.**
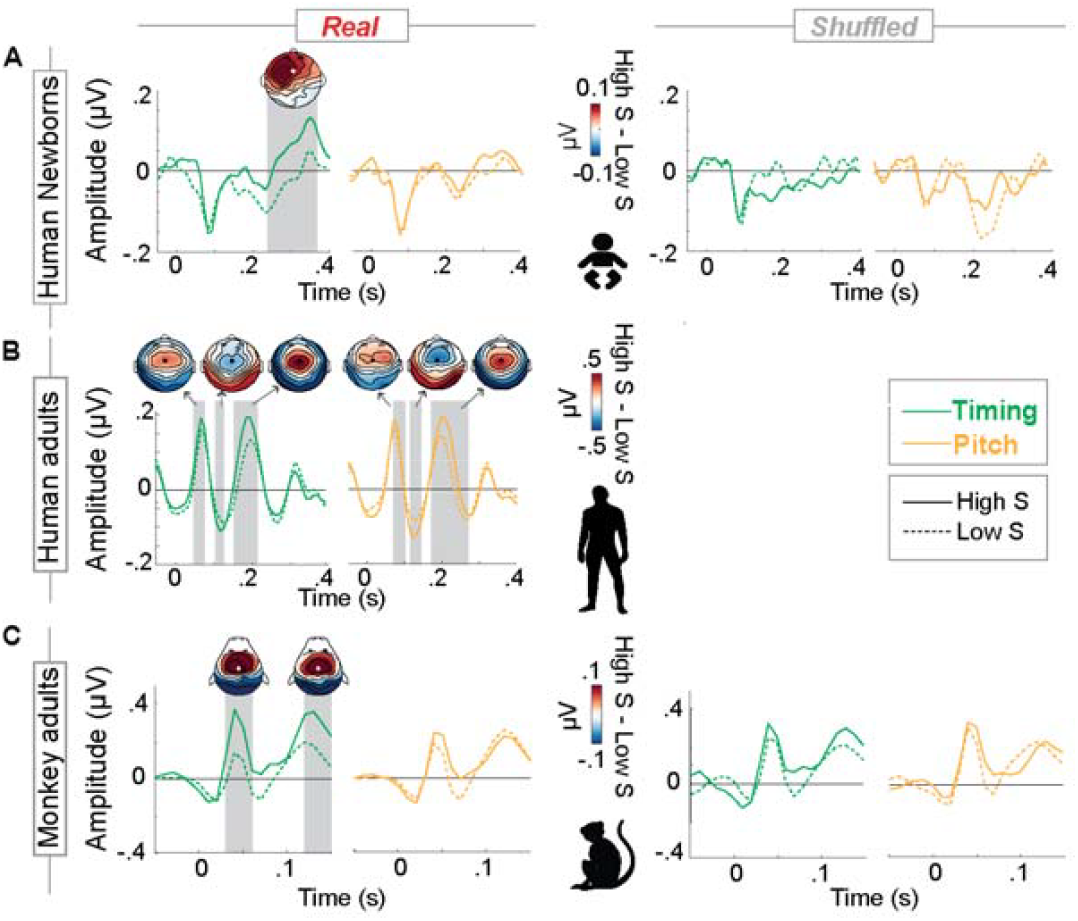
Modulation of auditory event-related potentials as a function of surprise. A) Human newborns. Group-average (n=49) ERPs (electrode Fz) evoked by notes carrying relatively high vs low St (green) and Sp (yellow) as a proxy of surprise (High S vs. Low S, respectively) for real and shuffled music locked to the note onset (0 s). The amplitude of the P2 component was higher in response to notes carrying relatively high vs low temporal surprise (green) (from +.24 to +.37 s) for real but not for shuffled music. No effect of pitch-related surprise was found. Grey windows highlight significant differences between low vs. high surprise responses (cluster-corrected permutation tests over time across all electrodes). Topographies illustrate the amplitude difference between conditions in the time windows of identified clusters. **B) Human adults**. To assist the comparison with results of previous studies, we plotted group-average (n=20) ERPs (electrode FCz) recorded from human adults listening to the same stimuli as the infants (reanalysis of data from (46)). Note that no shuffled stimuli were presented in this study. The amplitude of the P1-N1-P2 components was higher in response to notes associated with high than those with low temporal surprise (P1: from +.05 to +.07 s; N1: from +.11 to +.12 s; P2: from +.16 to +.21 s). Notably, human adults also exhibited sensitivity to pitch-related surprise, as indicated by enhanced P1-N1-P2 in response to notes carrying relatively high vs low surprise in pitch (Sp, yellow) (P1: from +.07 to +.09 s; N1: from +.12 to +.14 s; P2: from +.17 to +.27 s). **C) Adult Rhesus monkeys**. Group-average (n=2) ERPs (electrode FCz) recorded from Rhesus Monkeys listening to the same stimuli as the infants (reanalysis of data from (18)). The amplitude of the P1 and P2 components (P1: from +.03 to +.06 s; P2 from +.12 to +.15 s, frontal electrodes) was higher in response to notes associated with high than those with low temporal surprise for real, but not for shuffled music. Similarly to newborns, pitch-related surprise did not yield any significant modulation of the EEG amplitude.

Notably, the amplitude of the two positive-going deflections was enhanced in response to temporally unexpected (High S) compared to expected (Low S) notes, reaching significance in the second peak (from +.24 to +.37 s). This was observed for real but not for shuffled music. Conversely, no amplitude modulation was evoked by notes with unexpected pitch. These results fully align with the TRF results, confirming that babies generate expectations based on the rhythmic rather than melodic structure of the musical stimuli. They further provide insights into a neurophysiological response, specifically a late EEG positivity, whose amplitude varies as a function of timing-but not pitch-related surprise. Together, these results warrant comparison with findings from previous studies that exposed human adults (Fig. 3B) and monkeys (Fig. 3C) to the same stimuli (18, 46) to be further elaborated upon in the Discussion.

## Discussion

We employed continuous music stimuli with a rich melodic and rhythmic “alphabet” as a testbed for investigating the neurophysiology of music encoding in newborns. We demonstrated the feasibility of using ecologically valid complex stimuli, such as music, to examine different levels of auditory processing at birth. By determining how newborns use statistical regularities in melodic and rhythmic information to process music, our findings provide key contributions to understanding auditory development and its built-in biological constraints. Specifically, while rhythmic statistical regularities embedded in musical stimuli are neurally encoded already at birth, pitch-based information does not receive the same depth of processing, whether at low or high levels of encoding. This suggests that rhythmic and melodic sensitivities do not emerge in parallel in humans, with rhythm developing earlier than melody.

TRF analyses revealed high inter-individual variability in neural tracking of music (Fig 2B left panel, see also Jessen et al., 2021), likely stemming from the high variability in waveform’s morphology and latency of newborns’ auditory ERPs (54) and compatible with the notion that TRFs capture ERP-like responses (49). This variability could be explained by differences in sleep stages during testing, especially considering that sleep states modulate neural responsiveness (55) and, consequently, the individual infants’ capacity to encode continuous sound stimuli. Other potential sources of variability could be gestational age (56) or prenatal exposure to music (57). While we could not manipulate gestational age and/or prenatal exposure in the current study, future research should aim to systematically manipulate these factors under the hypothesis that greater neural maturation and/or musical exposure would lead to stronger neural encoding of musical expectations.

We showed that newborns track note-by-note predictability in real but not shuffled music. Importantly, the rhythmic, not the melodic, aspect of sound sequences drove this effect. This shows that newborns extract statistical regularities from structured contexts (real condition) – likely the rhythmic relationship between non-adjacent timing intervals – to predict upcoming events in the sequence. Conversely, musical expectations were reduced when regularities were weak or absent in random contexts (shuffled condition). Notably, local temporal information – the latency difference between two adjacent notes – was encoded while listening both to real and shuffled music, independently of whether high-order structural patterns were preserved. These findings align with the idea that tracking event predictability relies on the ability to extract and represent structural information from the past context (58, 59). The reduced response in the shuffled condition reflects a down-weighting of such predictability-related response when the inferred stability, or precision, of the sensory input is low and the present information does not conform with the past (60, 61).

This finding also brings novel evidence to our understanding of human rhythmic abilities present at birth. While rhythmic skills, such as sensitivity to isochrony and beat periodicity, are well-documented in infants, at 5 months (62), at birth (40), and even in preterm babies (39), evidence regarding sensitivity to context-based probabilistic expectations remains elusive (41). Here, we offer positive evidence. Using J. S. Bach’s compositions with a variable range of IOIs, we show that newborns are not merely tracking isochrony and periodic patterns. They also process a higher-level feature, namely the probability of when the next event will occur based on a range of past different IOIs. This capacity in infants might build upon the well-documented sensitivity to isochrony and periodicity: in other words, an isochronous or periodic representation of a sequence might provide a temporal grid of predictable sound events, like a scaffold facilitating the segmentation and organization of more complex temporal and/or spectral patterns (63). Such precocious rhythmic abilities might start and be significantly stimulated during gestation, given the prominence of biological rhythms in the fetal sensory environment. This includes auditory stimulations (e.g., the mother’s heartbeat, (64)), as well as vestibular stimulations (e.g., associated with the regular pace of maternal gait (65–67)). We speculate that this foundational sensitivity to rhythm might be key in the early development of cognition, not only as a precursor to higher-order statistical learning but also as a mechanism for adapting and organizing behavior in time. In support of this idea, newborns can partially adapt spontaneous rhythmical behaviors, such as sucking, to external stimuli (68); rhythmical rocking interventions on preterm infants improve orienting responses (69); and vestibular stimulation on preterm babies increases adaptive breathing response, vital for organizing structured behaviors, such as feeding, early vocalizations, and interactions (70).

As opposed to rhythm, we found no evidence for neural encoding of local pitch changes (IPI) or pitch-based probabilistic expectations in either of the music conditions. Whilst the lack of statistical significance for pitch-based probabilistic expectations could have been expected, the same for the tracking of local pitch changes is at first surprising, given previous evidence in newborns (42–45). However, it should be noted that IPI neural tracking approached significance when compared to zero in the real condition, and our study, compared with previous ones, used more complex, richer stimuli. Previous work (42–45) used sequences with no rhythmic variations (constant IOI), importantly, with an alphabet of a few pitches and IPI and highly infrequent, large pitch deviations. Across real and shuffled melodies, our musical stimuli instead were characterized by sequences of multiple pitches (N = 38, mean height = 73.04 ± 6.32; range: 55-93, in MIDI notation), IPIs (N = 29, mean interval size = 4.66 ± .4.12; range: 0-28) and IOIs (N = 30, mean interval size = .19 ± .13; range: .03-2.6) presented at varying tempi and different rhythms. Moreover, evidence of large variability and hardly detectable responses across neonates was also reported in previous work despite using temporally constant sequences and large spectral deviations (54). Overall, the absence of IPI tracking in our sample aligns with previous conclusions that reliable pitch neural tracking in newborns requires clear-cut pitch-change manipulations (54). This is likely related to immature frequency-specific pathways and coarse frequency tuning at birth (71), as well as to immature temporal resolution of different tones (see evidence from 6 m.o. infants (72)). It could also be a consequence of a gating effect of sleep on pitch-related features (see next paragraph). This limited sensitivity to deviant events suggests a rudimentary encoding of pitch relations in newborns compared to adults and possibly a longer developmental trajectory of pitch resolution compared to temporal resolution. According to this view, newborns’ sensitivity to pitch would be far from sufficient for appreciating musical pitch regularities, which likely emerge through maturation and enculturation.

We acknowledge that we cannot determine here whether the absence of significant melodic neural tracking in newborns reflects the true absence of a capacity that emerges through developmental refinement (as just discussed), or it is due to sleep-related suppression exacerbated by inherent pitch properties of a musical stimulus (pitch cues in other stimuli e.g. voice intonation, might be more salient to newborns). The lack of significant pitch-related (as opposed to rhythm-related) neural responses could reflect an adaptive prioritization of salient sensory processing during sleep: timing might be favored over pitch because it is more salient, relevant for attention, and potentially linked to survival-relevant cues (73). This would align with EEG studies on adults, suggesting that in-sleep perception and learning might be restricted to simple salient information (74). For example, higher-level auditory information is more suppressed than lower sensory one – meter-related more than meter-unrelated frequencies in music (75) and word/phrase more than syllable rates in speech (76). It is also possible that the greater complexity of pitch over timing information in music (Fig. 1B) made the pitch dimension too difficult for effective processing in newborns. Future research should investigate whether melodic processing is modulated by sleep in newborns and whether it is similarly underweighted in sleeping adults. This would clarify whether this effect is truly lacking in newborns or is a consequence of how pitch-related information is processed during sleep.

From a phylogenetic perspective, the prominent perceptual role of rhythm observed in early phases of human ontogeny might piggyback on a more ancestral phylogenetically conserved sensitivity to rhythm (rather than melody) within the primate lineage. The ERP analysis showed greater amplitude of P1-P2 components to temporally unexpected than expected notes but no modulation in the pitch dimension (Fig. 3A). Given their similar broad frontal topography, these two peaks may reflect a single positivity with a similar underlying generator (as also discussed in (51, 55)). They might also represent precursors to the adults’ P1 and P2 components (Fig. 3B), possibly involving frontotemporal areas for sensory predictive processing and memory-based sequential integration (77– 79). Interestingly, the P1-P2 responses of both monkeys and human adults listening to the same stimuli presented to the newborns were also modulated by temporal surprise (18, see the re-analyses in Fig. 3B-C). Thus, the similarity in cortical responses to temporal surprise across groups suggests rhythm as a primary perceptual cue in auditory sequence tracking. This does not necessarily imply that humans and monkeys generate rhythmic expectations through the very same neural mechanism, even if the cortical responses are similarly modulated by temporally unexpected events. For instance, these responses might rely on different anticipatory mechanisms sensitive to different rhythmic features— ranging from periodicity and local temporal changes to statistical and hierarchical structures (80, 81). Comparing distinct rhythmic computational models across phylogenetically close groups and as a function of exposure might shed light on the biological basis and evolutionary history of these different rhythmic capacities.

Regarding melodic expectations, the lack of significant melodic tracking observed in both human newborns and musically naïve monkeys (as opposed to human adults, Fig. 3) suggests that this ability is likely to be up-weighted with maturation and/or exposure to pitch-varying signals, such as speech and music. This leaves open the hypothesis that melodic sensitivity may not have emerged only in humans within the primate lineage but could potentially develop in other non-human primates given sufficient exposure to music during critical (and perhaps also non-critical) periods. Testing this hypothesis across species could shed light on the role of experience in shaping the relative weighting of pitch- and timing-based expectations in auditory processing.

Overall, this study provides neurophysiological evidence that tracking rhythmic statistical regularities is a capacity present at birth, whilst melodic tracking is not, at least for naturalistic musical stimuli such as the ones we used here. Future research should determine when this predominance of rhythm over melody changes across the first year of life, eventually reaching the stage observed for human adults, where melodic and rhythmic tracking have similar weights. This balancing process could parallel the onset of language development and social interaction (6-12 months), coinciding with an increasing awareness of speech as a key communicative tool (82).

## Methods

### Participants

64 healthy full-term newborn infants (0–2 days of age, 30 male, APGAR score 9/10 or above) were tested at the Department of Obstetrics-Genecology, Szent Imre Hospital, Budapest. EEG data from 6 babies were corrupted, and 9 other babies did not complete the experiment. As a result, these data were not analyzed, leaving a total sample size of 49. The analyzed babies had a mean gestational age of 40 weeks (SD = 7.1 days) and a mean birthweight of 3468.1 g (SD = 398.2 g). All newborns had normal hearing and passed the Brainstem Evoked Response Audiometry (BERA) test. Informed consent was obtained from at least one parent, and the infant’s mother could opt to be present during the recording. The study fully complied with the World Medical Association Helsinki Declaration and all applicable national laws and was approved by the Hungarian Medical Research Council, Committee of Scientific and Research Ethics (ETT TUKEB).

### Stimuli

The stimuli consisted of 14 monophonic piano melodies used in (Bianco et al., 2024; details in Table S1): 10 melodies (Real music) composed by Johann Sebastian Bach (previously also used in Di Liberto et al., and accessible here http://www.jsbach.net/) and 4 control melodies (Shuffled music) created by disrupting the pitch order and timing regularities of four of the original melodies (see below). The length of the melodies varied (average duration = 158.07 s ± 24.06), and the tempo ranged from 47 to 140 bpm (average tempo = 106.5 bpm ± 34.7). The four shuffled melodies were derived from four of the real melodies, specifically selected to represent those with the highest (melodies 05 and 08) and lowest (melodies 01 and 10) temporal-onset mean surprise. This selection was motivated by evidence suggesting that music with higher timing surprise elicits stronger brain responses in humans, aiming to balance these effects across both real and shuffled music. The shuffled melodies were matched to the real melodies in terms of pitch content, average note duration, and inter-onset intervals (IOIs), but their structure was disrupted in two key musical dimensions. Pitch regularities were altered by reordering the temporal sequence of the original notes. Rhythmic patterns were disrupted by creating a new set of inter-onset intervals (IOIs) drawn from a Gaussian distribution centered around the original mean IOI, with an added variation based on the difference between the mean and the minimum IOI. These randomly generated IOIs were then adjusted in MuseScore software (version 3.3.4.24412, https://musescore.org) to align with 16th-note quantization, preserving integer ratios. In MuseScore, the MIDI velocity (which correlates to note loudness) was standardized to a constant value of 100, and piano sound waveforms were synthesized with a 44,100 Hz sampling rate. Each melody was preceded and followed by a beep (800 Hz pure tone, linearly ramped with a 5 ms fade-in and fade-out) and a 5-second silence, following the structure: beep-silence-music-silence-beep. The resulting audio files were converted to mono and amplitude-normalized by dividing by the standard deviation using Matlab (R2019, The MathWorks, Natick, MA, USA).

### Information Dynamics of Music Model

Stimuli were analyzed using the Information Dynamics of Music (IDyOM) model (https://www.marcus-pearce.com/idyom/), which predicts note-by-note unexpectedness (surprise) and uncertainty (entropy). IDyOM is a variable-order Markov model that learns statistical patterns from musical sequences. It generates probability distributions for each new note based on prior context and outputs surprise (S) and entropy (E) over time. Surprise measures the unexpectedness of an event at time ‘t’ once it has occurred. Entropy reflects the uncertainty about the event at ‘t’ before it occurs based on the probability distribution of all potential notes considering all observations past the event at ‘t’. The model incorporates both short-term and long-term contexts, with the short-term model trained on the current sequence and the long-term model on prior musical exposure. To simulate the statistical knowledge that the newborns would acquire through mere exposure to the stimuli, predictions were derived from a combination of short and long-term models, with the latter being trained only on the stimuli used in the experiment, i.e. via resampling (10-fold cross-validation) (in IDyOM terminology: no pretraining, ‘‘both+’’ model configuration). IDyOM can account for many aspects of music, but here, we focused on two key dimensions that best describe piano monophonic melodies: pitch and timing. To this end, time series representing pitch and inter-onset interval ratios (using separate ‘cpitch’ and ‘ioi-ratio’ IDyOM viewpoints) were analyzed independently by IDyOM to calculate note-by-note surprise (S) and entropy (E) for both pitch (Sp, Ep) and timing (St, Et). These were then combined to determine the joint (sum) probability for each note (S, E). To simulate the long-term statistical knowledge of music possibly acquired by babies in the womb, we run control analyses by using S and E estimates derived by an IDyOM model pre-trained on a large corpus of music (comprising 152 Canadian folk songs, 566 German folk songs from the Essen folk song collection, and 185 J. S. Bach chorale melodies (as in previous applications (18, 46, 83)). These analyses using the pretrained IDyOM model configuration led to a similar pattern of results (Fig. S1). This observation suggests that Bach’s selected musical pieces may already contain enough regularities for the model to learn directly from the stimulus set, making pretraining on a larger corpus redundant.

### Procedure

As a common procedure in EEG studies in newborns, babies were asleep during the EEG recording and stimulus presentation. Stimuli were presented using a Maya 22 USB external soundcard and ER-2 Insert Earphones (Etymotic Research Inc., Elk Grove Village, IL, USA) placed into the infants’ ears via ER-2 Foam Infant Ear-tips. The melodies were presented at a comfortable intensity (about 70 dB SPL). Two sets of the 14 melodies were presented in a randomized order within each set, ensuring that each baby listened to each melody at least once, with some melodies being heard twice. The presentation was implemented in Matlab (R2014, The MathWorks, Natick, MA, USA) and Psychtoolbox (version 3.0.14). EEG was recorded throughout the stimulus presentation. The inter-stimulus interval between melodies (ISI, offset to onset) was 900–1300 ms (random with even distribution, 1 ms step). The experiment took 45 min overall, including both preparation and stimulation.

### Data recording and pre-processing

An ActiChamp Plus amplifier with a 64-channel sponge-based electrode system (saltwater sponges and passive Ag/AgCl electrodes, R-Net,) and Brain-Vision Recorder were employed to record EEG (Brain Products GmbH, Gilching, Germany). The sampling rate was 500 Hz with a 100 Hz online low-pass filter applied. Electrodes were placed according to the International 10/10 system. The Cz channel served as the reference electrode while the ground electrode was placed on the midline of the forehead. During the recording, impedances were kept below 50 kΩ.

Data were pre-processed and analyzed in Matlab R2019. For the analysis, we applied a fully data-driven pipeline for preprocessing EEG data, combining open-access denoising algorithms, similar to previous studies dealing with noisy EEG recordings (18, 52). The analysis used Fieldtrip (84) and EEGLAB toolboxes (http://sccn.ucsd.edu/). The continuous EEG data were bandpass filtered between 1 and 30 Hz (Butterworth filter, zero-phase, order 3), down-sampled to 100 Hz, and segmented into epochs from the onset to the offset of each melody, separately. Before re-referencing the data to the average of a set of electrodes (‘F9’, ‘F10’, ‘P9’, ‘P10’, ‘Iz’), faulty or noisy electrodes were temporarily discarded to prevent noise contamination across electrodes. Specifically, for each electrode, the mean, standard deviation, and peak-to-peak values were calculated across time within each trial. If any of these values deviated by more than 2.75 standard deviations from the mean of other electrodes, the electrode was flagged as noisy/faulty. This process was repeated until a distribution without outliers was obtained. The data were then further denoised in EEGLAB using the Artefact Subspace Reconstruction (ASR) algorithm (85) (threshold value 5 previously validated for both adult human and monkey EEG data (18)). Eye-movement artifacts were corrected using the ICLabel algorithm in EEGLAB. After performing independent component analysis (ICA) with EEGLAB’s ‘runica’ function, independent components labeled by ICLabel as ‘eye movements’ (with > 90% likelihood) were rejected. Subsequently, electrodes that were initially excluded (due to being faulty or noisy) were interpolated by replacing their voltage with the average voltage of the (preprocessed) neighboring electrodes (18 mm distance, including 8 electrodes on average). If, following the above pre-processing, noisy electrodes were still automatically identified, the interpolation step was repeated (the number of such iterations varied between 1 and 2).

### TRF Analysis

We employed Temporal Response Functions (TRF) to model EEG responses to the continuous acoustic and musical features of the presented stimuli using the mTRF Matlab toolbox (48). Each stimulus feature (as listed below) was normalized across time for each melody, ensuring that the root mean square of each feature was 1. A forward model was run to predict the ongoing EEG response from the stimulus features, with a time lag window of -50 to +400 ms to capture EEG fluctuations related to changes in the stimulus. This time window was sufficiently large to encapsulate well-known ERP-like modulations of EEG signals that are known to drive the variance modeled by TRF. Ridge regression was used to prevent overfitting (lambda range: 10^−4^ to 10^2^). TRFs were fitted to all melodies (pooled real and shuffled melodies) using leave-one-melody-out cross-validation, and the EEG time course of the left-out melody was predicted. Note that the correlation values are typically calculated between EEG signals and their predictions by considering single-participant EEG signals, which might carry much noise (especially if recorded from newborns). As such, EEG prediction correlations are variable between participants largely due to the variable SNR of the EEG signal across participants (as every prediction is correlated with a different EEG signal). To overcome this issue, we averaged all participants’ EEG timeseries data to form a single EEG ‘super-subject’ data timeseries, which we refer to as ‘ground-truth EEG’. Then, per each participant, melody, and electrode, prediction accuracy was quantified by calculating Pearson’s correlation between the predicted and ground-truth EEG data.

We tested the contribution of high-level probabilistic musical expectations to the predicted EEG in addition to that of the low-level acoustic features in both the timing and pitch dimensions. Feature selection was based on the approach used in (18), but we additionally tested for the contribution of local changes in timing and pitch, such as inter-onset- and inter-pitch-intervals (IOI, IPI, measured in ms and absolute number of semitones, respectively), as these features are often correlated with Surprise values in real music (86) (see Fig. 1C). Note, however, that analyses excluding these features led to similar conclusions as derived from the main analyses. Therefore, we run a full model including low-level acoustic features (acoustic onset, spectral flux, as well as IOI and IPI) and high-level probabilistic musical features with impulses at the note onsets but whose amplitudes are set to the pitch and onset surprise and entropy values from IDyOM (Surprise pitch, Surprise timing and Entropy pitch, Entropy timing – Sp, St and Ep, Et). We then assessed the unique neural encoding of a single or a set of stimulus features by subtracting the prediction accuracy of several reduced models from the full model (containing all features). The reduced models had the same dimensionality as the full model, but the feature/s of interest was/were randomized in time whilst preserving the onset times. We tested five reduced models: 1) A probabilistic music model where the high-level features (St and Et and Sp and Ep) were randomized to assess the overall effect of adding surprise and entropy estimates to low-level features to the neural tracking; 2) A probabilistic timing model with randomized St and Et; 3) A probabilistic pitch model with randomized Sp and Ep; 4) A local timing model with randomized IOI; 5) A local pitch model with randomized IPI. To compare across the different models, for each participant, condition, and TRFmodel, Δ*r* values were averaged across 25% of channels with the highest prediction accuracy in the full model and across the real and the shuffled condition. These values were then entered into linear mixed-effects regressions.

### Statistical analysis

Statistical analyses were run in R (version 4.1.3, 2022-03-10) and included non-parametric tests or linear mixed-effects models (lme4 package). All models included Random Effects of Infants and Melodies (IDs 1-14). The Fixed effects included Condition (real/shuffled) and TRFmodel (depending on the comparison; see results). Statistical significance was evaluated by likelihood-ratio tests (χ^2^) conducted using the ‘anova’ function (stats package). Follow-up contrasts were conducted using the ‘emmeans’ package and the Tukey method to account for the increased risk of type I error resulting from multiple comparisons. Adjusted p-values were calculated to determine significant differences between conditions. A significance level of a = 0.05 was used. All linear mixed-effects models (LMMs) report fixed-effect estimates (b) along with their standard errors (SE) and t-values, with degrees of freedom estimated via Satterthwaite approximation when applicable.

When a direct test of differences was needed, the non-parametric Wilcoxon signed-rank test was used. For these, results are reported as W-values, indicating the sum of ranks of signed differences.

### ERP Analysis

Event-related potential (ERP) analyses were performed by segmenting the EEG data into 600 ms epochs, beginning 100 ms before the onset of each note and ending 500 ms after the onset. Epochs were baseline corrected using a 50 ms window before the note onsets, and trials that deviated from the mean by more than 2.5 the average standard deviation were rejected (3.45 ± 1.12 % of the trials per subject). To assess ERP modulation based on note surprise, we selected the notes with the highest and lowest 20% surprise (high S and low S) values, separately for each melody, as assessed by the IDyOM. Per each subject, epochs were trimmed to a window of -50 +400 ms relative to note onset and averaged by high/low S condition, separately for real and shuffled melodies. Cluster-based permutation testing (87) was used to account for multiple comparisons across adjacent time points and electrodes. Clusters of adjacent timepoints and neighboring electrodes (at least three) associated with significant (p-values < 0.025) differences across conditions were formed. A cluster-level threshold of p < 0.05 was applied to the t-statistic and the Monte Carlo method (1000 iterations) was used to estimate the null distribution of this statistic. To assist comparability with the previous work, we re-analyzed the EEG data recorded from human (46) and monkey (18) adults following the same pipeline described here (both datasets are open source). Note that for the monkey data, as in the original work, clusters were identified separately for each animal (across 22 sessions) and considered significant only when exhibited by both animals (conjunction analysis).

## Data Availability Statement

The data reported in this manuscript will be made available upon publication in the following repository: https://doi.org/10.48557/GMOFYE. The codes used for analyses and figures are available in the following GitHub repository: https://github.com/robilobi/humannewborns_eeg_music.

## Acknowledgements and funding sources

R.B. is funded by the European Union (MSCA, PHYLOMUSIC, 101064334). G.N. and F.B. are funded by the European Research Council (ERC, MUSICOM, 948186). T.N. is funded by the European Union (MSCA, SYNCON, 101105726). BT, GPH, and IW are funded by the Hungarian National Research Development and Innovation Office (ANN131305, FK139135, and K147135, respectively).

## Authors’ contribution

Roberta Bianco: Conceptualization, Methodology, Formal Analysis, Investigation, Data curation, Visualization, Writing Original draft, Writing - Review & Editing, Funding acquisition; Brigitta Tóth: Investigation, Resources, Writing - Review & Editing, Funding acquisition, Project administration; Félix Bigand: Methodology, Writing – Review & Editing; Trinh Nguyen: Methodology, Writing – Review & Editing; Gábor Háden: Investigation, Writing - Review & Editing; István Sziller: Investigation, Data curation. István Winkler: Writing - Review & Editing, Funding acquisition; Giacomo Novembre: Conceptualization, Methodology, Writing Original draft, Writing Review & Editing, Funding acquisition, Supervision.

## Competing interests

The authors declare no competing interests.

## Additional information

**Supplementary information** This manuscript contains supplementary materials (see supplementary document).

## Supplementary material

**Figure S1.**
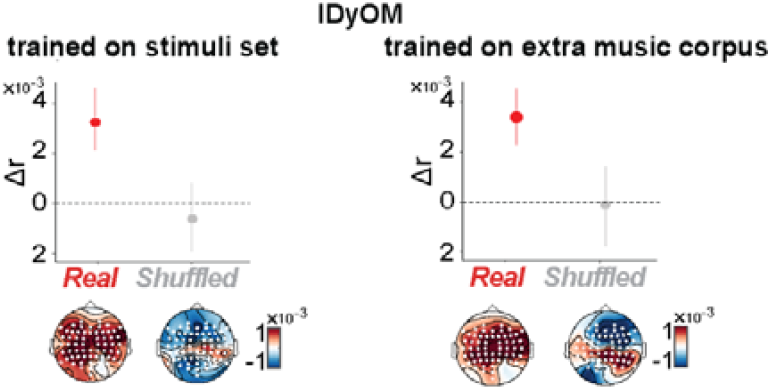
Control analyses. No effects of IDyOM statistical knowledge on EEG prediction accuracy. We compared the effect of deriving surprise estimates by training IDyOM on either the experimental stimuli alone (left panel) or on the experimental stimuli, as well as an additional corpus of western tonal music (right panel). For each panel, we plot the difference in EEG prediction accuracy (Δr) between the full and the reduced models (randomizing St and Et and Sp and Ep). Dots represent the grand-average mean Δr computed across all channels and melodies (left top panel, with associated topographical maps) for real (red) and shuffled (grey) music. Error bars represent bootstrapped 95% CI. The absence of differences in predicting neural responses between pre-trained and non-pre-trained model configurations suggests that incorporating pretraining to estimate surprise and entropy values does not enhance our ability to predict EEG data. This may be due to the high correlation between the estimates derived from the two IDyOM configurations, leading to similar EEG predictive power. Additionally, it may indicate that Bach’s music contains sufficient rules and statistical regularities, allowing the model to learn these directly from the stimuli set, rendering pretraining on the large music corpus redundant.

**Table S1.** Experimental stimuli.

| Melody # | Title | IOI (s)<br>(mean, SD, range) | Pitch (MIDI) (mean, SD, range) | Melody Duration (s) | #Notes |
| --- | --- | --- | --- | --- | --- |
| 01 | BWV1013 allemande | $0.15 \pm 0.04$<br>(0.75:1.35) | $75.3 \pm 5.5$<br>(62:93) | 154 | 1021 |
| 02 | BWV1013 corrente | $0.17 \pm 0.08$<br>(0.75:0.9) | $74.9 \pm 5.3$<br>(62:91) | 150 | 890 |
| 03 | BWV1013 sarabande | $0.38 \pm 0.25$<br>(0.21:2.57) | $76.7 \pm 5.7$<br>(62:92) | 116 | 300 |
| 04 | BWV1013 bourrée angloise | $0.25 \pm 0.13$<br>(0.19:1.12) | $73.8 \pm 4.9$<br>(60:86) | 132 | 528 |
| 05 | BWV 1002 allemande | $0.38 \pm 0.18$<br>(0.07:1.5) | $73.4 \pm 6.6$<br>(57:86) | 170 | 619 |
| 06 | BWV 1002 gigue | $0.3 \pm 0.13$<br>(0.16:2.55) | $70.6 \pm 5.6$<br>(56:86) | 161 | 539 |
| 07 | BWV 1002 gavotte | $0.13 \pm 0.04$<br>(0.12:0.5) | $72.0 \pm 6.2$<br>(55:86) | 179 | 1351 |
| 08 | BWV 1006 loure | $0.38 \pm 0.19$<br>(0.12:1.5) | $77.6 \pm 4.4$<br>(64:85) | 130 | 337 |
| 09 | BWV 1006 gavotte | $0.27 \pm 0.15$<br>(0.11:1.29) | $75.3 \pm 4.9$<br>(59:86) | 175 | 641 |
| 10 | BWV 1001 presto | $0.12 \pm 0.03$<br>(0.12:0.72) | $70.0 \pm 6.1$<br>(55:86) | 195 | 1603 |
| s_01 | Shuffled o_01 | $0.15 \pm 0.05$<br>(0.04:0.31) | $75.3 \pm 5.5$<br>(62:93) | 152 | 1021 |
| s_05 | Shuffled o_05 | $0.27 \pm 0.134$<br>(0.06:0.9) | $73.4 \pm 6.6$<br>(57:86) | 166 | 619 |
| s_08 | Shuffled o_08 | $0.39 \pm 0.18$<br>(0.05:1.15) | $77.5 \pm 4.4$<br>(64:85) | 133 | 337 |
| s_10 | Shuffled o_10 | $0.12 \pm 0.02$<br>(0.06:0.18) | $70.0 \pm 6.1$<br>(55:86) | 193 | 1603 |

## Notes

### Competing Interest Statement

The authors have declared no competing interest.

